# MixMir: microRNA motif discovery from gene expression data using mixed linear models

**DOI:** 10.1101/004010

**Authors:** Liyang Diao, Antoine Marcais, Scott Norton, Kevin C. Chen

**Affiliations:** BioMaPS Institute for Quantitative Biology and Department of Genetics, Rutgers, The State University of New Jersey, Piscataway, NJ 08854, USA; CIRI, International Center for Infectiology Research, Université de Lyon, Inserm, CNRS, Ecole Normale Supérieure, Lyon, France; Department of Mathematics and Department of Molecular and Cell Biology, University of Connecticut, Storrs, CT 06269, USA

## Abstract

microRNAs (miRNAs) are a class of ∼22nt non-coding RNAs that potentially regulate over 60% of human protein-coding genes. miRNA activity is highly specific, differing between cell types, developmental stages and environmental conditions, so the identification of active miRNAs in a given sample is of great interest. Here we present a novel computational approach for analyzing both mRNA sequence and gene expression data, called MixMir. Our method corrects for 3′ UTR background sequence similarity between transcripts, which is known to correlate with mRNA transcript abundance. We demonstrate that after accounting for *k*mer sequence similarities in 3’ UTRs, a statistical linear model based on motif presence/absence can effectively discover active miRNAs in a sample. MixMir utilizes fast software implementations for solving mixed linear models which are widely-used in genome-wide association studies (GWAS). Essentially we use 3’ UTR sequence similarity in place of population cryptic relatedness in the GWAS problem. Compared to similar methods such as miReduce, Sylamer and cWords, we found that MixMir performed better at discovering true miRNA motifs in three mouse Dicer knockout experiments from different tissues, two of which were collected by our group. We confirmed these results on protein and mRNA expression data obtained from miRNA transfection experiments in human cell lines. MixMir can be freely downloaded from https://github.com/ldiao/MixMir.

## INTRODUCTION

microRNAs (miRNAs) are small (∼22nt) non-coding RNAs that post-transcriptionally regulate the expression of protein-coding genes (1). Their impact on gene regulation is the subject of intense study, with over 60% of all human genes estimated to be regulated by miRNAs (2) and some miRNAs potentially regulating hundreds of genes (3,4). Thus computational prediction of active miRNAs and their targets from gene expression data in a particular cellular context is of significant interest, leading to the development of a number of algorithms that analyze miRNAs jointly in the context of sequence and gene expression (5–9).

Animal miRNAs can bind to their targets in a variety of ways, centering on a 6nt region at the 5′ end of the mature miRNA (bases 2-7) called the “seed” region (4). Most computational target prediction methods make use of exact Watson-Crick pairing of the seed region, as well as other features such as evolutionary conservation (10,11), co-occurrence of miRNA binding sites (12) and RNA-binding protein binding sites (13), mRNA, miRNA, and Argonaute expression levels (5–7,9,13), 3’ UTR sequence composition and other mRNA sequence features (5,6,12,13), and protein interaction data (14). A commonly-used computer program for analyzing miRNAs from both gene expression and sequence data is miReduce (7), which is based on the REDUCE algorithm for predicting transcription factor motifs (15). Two other published programs, Sylamer (5) and cWords (6), also solve the same problem while explicitly correcting for background sequence composition. The context of the miRNA binding site is known to affect binding efficacy (12), either through mRNA secondary structure or by providing binding sites for other post-transcriptional regulators. Background sequence similarity is also correlated with paralogy and therefore similarity between transcriptional programs. Note that it is not trivial in mammalian genomes to identify promoter or enhancer regions in order to directly correct for transcriptional control of protein coding genes.

Here we present MixMir, a novel method for miRNA motif discovery which, like Sylamer and cWords, explicitly corrects for background sequence composition, but does so using a mixed linear model framework. In our MixMir implementation, we borrowed computational methods from the genome-wide association studies (GWAS) field for efficiently solving large systems of mixed linear model (MLM) equations. Our method uses the MLM to correct for similarities between 3′ UTR sequences, analogous to the way MLMs are used to correct for cryptic relatedness between individuals in GWAS, where such artifacts can lead to high false positive associations (16–18).

We demonstrate the utility of MixMir on a gene expression data set in mouse wildtype and Dicer knockout CD4+25- T-cells (hereafter called T conv cells). We found that MixMir performed better than miReduce, Sylamer, and cWords at finding miRNAs annotated in the miRBase database (19) and highly expressed miRNAs in this cell type. We confirmed our results on two other mouse Dicer KO experiments from embryonic stem cells and adrenal gland, and miRNA transfection experiments in human cell lines, both for quantitative proteomics data and microarray data (20). Thus we expect MixMir to be of practical use in helping experimentalists focus attention on the most important miRNAs in a given sample. Importantly, the most active miRNAs (i.e. those that play the biggest role in controlling mRNA expression in a particular cell type) are positively but imperfectly correlated with miRNA expression levels in the cell. Thus, it is not sufficient to simply assay miRNA expression levels in the cell to identify the most active miRNAs (see Discussion). Another application where miRNA motif finding is often used in the lab is simply to confirm that a transfection or knockdown experiment worked correctly.

More broadly, our study suggests that mixed linear models are a powerful tool for motif discovery and highlight the importance of correcting for cryptic similarity in background sequence composition, an observation which might be useful in other motif-finding problems as well. Our MixMir software is open source and freely available from https://github.com/ldiao/MixMir.

## MATERIAL AND METHODS

### Experimental methods for obtaining mRNA and microRNA expression data for Dicer KO T conv cells

Mice carrying a floxed Dicer allele in combination with CD4Cre transgene on a mixed C57BL/129 background (21) were maintained under specific pathogen-free conditions. Peripheral CD4+CD25- T cells were sorted on a FACS ARIA (Becton Dickinson) from 6-8 week-old mice and RNA extracted using RNAbee (AMSBio) according to the manufacturer’s instructions. 100 nanograms of RNA was used to interrogate the GeneChip Mouse Gene 1.0 ST Array (Affymetrix). We obtained log fold changes in gene expression for 24,601 mRNA transcripts between WT and Dicer KO mouse CD4+ CD25- T cells.

We obtained two data sets of miRNA expression for CD4+CD25- T cells from independent sources using different technologies. First, from the same cells from which we obtained our mRNA microarray expression data, we also obtained comparative miRNA expression data between CD4+CD25+ T-cells and CD4+25- T-cells from Cobb et al. (22), who studied the differences in miRNA expression profiles for the two types of cells. The authors performed a miRNA microarray analysis with probes for 173 miRNAs from miRBase. Of these, we take the top 20 differentially expressed to be true, “active” miRNAs, as reported by Cobb et al. in Figure 2.

To corroborate these results, we also used miRNA expression data from C57BL/6 mice determined by the nCounter miRNA expression assay kit (Nanostring Technologies), from Sommers et al. (23). The authors validated the Nanostring nCounter expression results with Exiqon microarrays and Taqman qRT-PCR assays.92 probes corresponding to 86 miRNAs in miRBase were evaluated. Of these, we took the top 21 highly expressed for experimental validation of our predictions, corresponding to the most highly expressed miRNAs presented by Sommers et al. in Figure S1.

### Experimental methods for obtaining mRNA expression data for Dicer KO mouse embryonic stem cells

The ES cells were derived and described in Nesterova et al. (23). ES cell lines were maintained on a feeder layer (mitomycin-inactivated primary mouse embryonic fibroblasts) in Dulbecco’s Modified Eagle Medium (DMEM) supplemented with 10% fetal calf serum (FCS, Autogen Bioclear), 7% Knockout Serum Replacement (KSR), 2 mM L-glutamine, 1× non-essential amino acids, 50 μM 2-mercaptoethanol, 50 μg/ml penicillin/streptomycin (all from Invitrogen) and LIF-conditioned medium, made in house, at a concentration equivalent to 1000 U/ml. Cells were grown at 37°C in a humid atmosphere with 5% CO2. Microarray methods were the same as for the T conv cells as described above. Sinkkonen et al. (24) showed that in mouse ES cells, transfection of miR-290 family miRNAs is able to rescue defects due to Dicer deficiency. This strongly suggests that the primary active miRNA in these cells belong to the miR-290 family.

### pSILAC and microarray data for miRNA transfection experiments

In addition to our Dicer KO data, we also tested MixMir on pSILAC and mRNA microarray data from miRNA transfection experiments from Selbach *et al*. (20). The authors performed transfections by synthetic miRNAs and mock transfections in human HeLa cells for miRNAs let-7b, miR-1, miR-155, miR-16 and miR-30a. The amount of protein synthesis was given by the log of the ratio of protein synthesized in the miRNA transfected cells divided by the mock transfection between 8hrs and 32hrs post transfection. Microarray analyses were performed with the Affymetrix Human Genome U133 Plus 2.0 chip.

We mapped the pSILAC Uniprot protein IDs to Refseq transcript IDs by downloading an ID mapping table from the Uniprot website. For the different transfection experiments, there were slightly different numbers of proteins with expression values, resulting in a range of the number of protein expression data points with corresponding 3′ UTR sequences from ∼3000 - 3600 across all the transfection experiments.

### mRNA and miRNA expression data for Dicer knockout mouse adrenal cortex cells

We also obtained mRNA and miRNA expression data for Dicer KO adrenal cortex tissue from mouse embryos at stages E15.5 and E16.5 from a study by Krill et al. (25). *Sf1-Cre* mice were crossed with mice carrying a floxed Dicer allele to produce *Sf1-Cre/Dicer^lox/lox^* mice. Embryos were harvested at E15.5 and E16.5 and the adrenals from each were collected. A total of 4 control and 4 Dicer KO biological replicates were obtained for each time point. Affymetrix Mouse 430 v2.0 gene expression arrays were used for hybridization. ABI miRNA OpenArray was used for miRNA expression analysis. These arrays are able to target 750 miRNAs. We took the top 25 highly expressed miRNAs for E15.5, the top 35 highly expressed miRNAs for E16.5, and the top 15 miRNAs which are found highly expressed at both time points, in line with Figure 4 and Table 1 in Krill et al. (25).

**Table 1.** AU content of motifs discovered by the different methods. Simple linear models and cWords2 returned motifs with very high AU content. Both MixMir and miReduce had substantially lower average AU content, closer to the background 3’ UTR base composition.

| Method | % AU in motif |
| --- | --- |
| LM Bin | 88.67 |
| miReduce | 43.33 |
| Sylamer | 58.33 |
| cWords2 | 81.67 |
| MixMir6 | 53.67 |

### Processing of the 3′ UTR and miRNA sequence data

We downloaded all 26,845 mouse RefSeq gene 3′ UTR sequences and 40,571 human 3′ UTR sequences from the UCSC Genome Browser (version mm10 and hg19, respectively) (26,27). We removed all 3’ UTRs of length less than 10nt and retained the longest isoform if there were multiple 3’UTR isoforms. In total, we were able to associate 17,988 unique UTR sequences to their microarray expression values for the mouse Dicer KO dataset, and 22,266 unique UTR sequences to their microarray expression values for each of the Selbach *et al*. miRNA transfection experiments.

We downloaded 1,908 mature mouse miRNA sequences corresponding to ∼1200 distinct 6mer seeds and 2,578 mature human miRNA sequences corresponding to ∼1500 distinct 6mer seeds from the miRBase database (release 20) (19).

### Linear regression model of miRNA targeting

A naive linear model formulation of miRNA targeting is to regress the log fold change in gene expression against the presence/absence of the miRNA motif in the 3′ UTR:

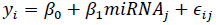

where *y_i_* is the log fold change in expression level of mRNA *i* and *miRNA_i,j_* is the presence/absence variable that indicates if the motif for miRNA *j* appears in the 3’ UTR of mRNA *i*, the *β* variables are constants and ε*_ij_* is a Gaussian error term. We applied this simple linear model for all potential miRNA seeds (i.e. the set of all 4096 hexamers) across all mRNAs. The null hypothesis is that there is no miRNA effect, i.e. *β*_1_ = 0. If the deviation of the inferred *β*_1_ from 0 is statistically significant, then we say that the presence of the motif for miRNA *j* is significant.

### Mixed linear model of miRNA targeting

Our mixed linear model builds on the simple linear model above by adding an additional random effect to account for pairwise background sequence similarity between 3′ UTRs. A random effect is a factor which can be modeled as being drawn from a probability distribution and is commonly used to model hierarchical structures in data (28). Here we take the set of all background *k*mer compositions of all 3′ UTRs as the distribution and consider each 3′ UTR to be a sample from it. Specifically, our mixed linear model is formulated as:

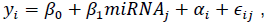

where *α_i_* is the random effect for the *i*th mRNA and *miRNA_i,j_* is the binary variable representing the presence or absence of the hexamer motif associated with miRNA *j* in individual *i*. Rewriting the above in matrix notation gives:

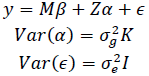

Here *y* represents the *T* × 1 vector of mRNA expression changes, where *T* is the number of mRNAs, M is a binary diagonal matrix indicating the presence or absence of a particular miRNA, i.e. *M_j,j_* = *miRNA_j,_* and Z is an incidence matrix (in our case *Z* = *I_T_*). The most important part of the model is *α*, a vector of random effects, which incorporates a constraint on the covariance matrix via the *T* × T relationship matrix *K*, which we set to be the pairwise relationship matrix between mRNAs, as discussed below. *ε* and *α* are multivariate Gaussians with mean 0. 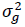 and 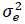 are called the variance components of the model.

We let *K_ij_* = *cor*(*kmer_i,_ kmer_j_*), the pairwise Pearson correlation for the fractional *k*mer counts between mRNA *i* and mRNA *j*. The fractional *k*mer counts are obtained by taking the *k*mer counts and scaling by the sum of the total number of motif counts:

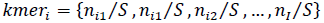

where *n_ik_* is the number of times the motif *m_k_* appears in the 3′ UTR of mRNA *i* and 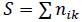 is the sum of the motif counts. We tested various *k*mer lengths in the construction of the relationship matrix in the range *k* = 2,…,6.

We used FaST-LMM v2.07 (29) to solve for the restricted maximum likelihood (REML) solution for the mixed linear models. It is an “exact” MLM solver in the sense that other fast solvers commonly used in GWAS often make simplifying assumptions specific to GWAS applications that are often not appropriate in the miRNA context. Briefly, FaST-LMM reparamaterizes the optimization problem in the mixed linear model to be a function of only a single parameter δ and then performs a spectral decomposition of the relatedness matrix once that can be used to test all motifs (or SNPs in the GWAS problem).

### Overview of the miReduce, Sylamer and cWords algorithms

miReduce takes the log fold change of gene expression between two conditions as input and outputs significant motifs and their associated miRNAs (7). The underlying algorithm is a forward stepwise linear regression, an iterative procedure where at each iteration the motif which minimizes the residual error in a simple linear model is selected and the residuals are taken as the new dependent variable for the following iteration. This continues until a significance threshold set by the user is exceeded. Forward stepwise linear regression procedures are known to suffer from many statistical problems, so the *p*-values from such procedures should be taken only as a general guideline (30). In this work we set a liberal cutoff of *p* = 0.50 for miReduce for the purpose of comparing the miReduce motifs with the linear models described above.

We ran Sylamer via its web-based implementation, Sylarray (5,31), because the Sylamer code did not collapse low complexity and redundant sequences, and preprocess 3′ UTR transcript variants which are implemented in Sylarray and are important components of the overall Sylamer method. Given a gene list of *N* genes ranked in descending order of differential gene expression, the Sylamer algorithm computes over- and under-representation of motifs in the top *T* genes vs. the remaining *N - T* in the list, as *T* is incremented in bin sizes of *b*. The hypergeometric distribution is used to determine the significance of the enrichment or depletion of a particular motif *m* in the top *T* genes, compared to the rest of the gene list, given that the total number of genes containing the motif in its 3′ UTR is *K_m_*. Sylamer corrects for background sequence composition by estimating expected motif counts based on the sequence composition of shorter motifs within each bin and using these values in place of *K_m_* (5). We use the default parameters for Sylarray, except we select the “all words” option, which searches the space of all possible 6-, 7-, and 8mer motifs instead of just those corresponding to known miRNAs. To obtain a full list of ranked motifs, we downloaded the enrichment table for each analysis and ranked motifs by their most significant *p*-value across all bins (see Sylamer manuscript for a full description of Sylamer defines statistical significance).

cWords (6) corrects for background sequence composition using a *k*th order Markov model. cWords estimates the expected probability of seeing a particular word based on the probability of seeing shorter words within the same UTR sequence, instead of looking through bins of motifs. The probability that a motif is enriched is calculated using the binomial distribution and the negative log of the probabilities is plotted to show enrichment across all ranked genes (genes ranked by increasing differential expression). Background enrichment values are computed as a sum of all such log probabilities, called the “running sum”, and enriched motifs are those that have statistically higher sums than the expected maximum value. We downloaded cWords from https://github.com/simras/cWords and ran it with word length = 6 and order of Markov background nucleotide model in the range 2-6. The Markov model in cWords can correct for kmers of length up to the motif length being analyzed. Thus, as we are analyzing hexamers, we can use this model to correct for background *k*mers with *k* ≤ 6. In principle higher values of k should give more correction but in practice the amount of data available for estimating the Markov model is limited by the length of the 3’ UTR sequences (see Results, Table S2). Thus we found that correcting for a background kmer of length *k* = 2 produced the best results which is entirely consistent with the authors’ recommendation to set k between 1 and 3.

Both Sylamer and cWords return results separated into those motifs being either enriched or depleted. For both methods we pooled the results to be more comparable with the other models, as we are looking for any significant motifs regardless of the direction of effect. We reserve the direction of the effect as an independent test of the accuracy of the methods later.

## RESULTS

### Comparison of different MixMir and cWords parameters on the T conv microarray data

We started by testing five different settings of the *k*mer length in MixMir on the mouse Dicer-knockout T conv microarray data set (Methods). MixMir uses a linear model of miRNA targeting, similar to previous models such as miReduce, but adds an additional relationship matrix in a mixed linear model framework to correct for background sequence composition of 3′ UTRs (Methods). The kmer length determines the construction of the relationship matrix in MixMir (Methods). We tested values of *k* from 2 to 6 and refer to these models as MixMir2 to MixMir6. For values of k above 6, the estimate of the correlation matrix became inaccurate because of the limited total amount of 3′ UTR sequence in the genome and the running time of the implementation was slow, so we did not consider higher values of k further (Discussion).

The value of *k* had a strong effect on the similarity of MixMir to the baseline linear model we tested (Methods), where similarity was defined by the Pearson correlation of the *p*-values of the motifs tested. Namely, the similarity of MixMir to the linear model dropped as *k* increased in MixMir. These patterns were similar if we computed the Pearson correlation of motif ranks instead of motif *p*-values (Table S1).

We compared percentile-percentile (PP) plots of all of the MixMir models to each other and to the linear model, to determine how skewed the *p*-values were across all motifs for each method (Figure 1). Under the null hypothesis of no association, we expect the curves to fall on the diagonal line *y* = *x*, so the presence of skew away from this line is indicative of false-positive associations. These plots clearly showed that the MixMir model which performed the most correction of the *p*-values was MixMir6. Note that this analysis implicitly assumes that there are relatively few highly active miRNAs in any particular cell type compared to the total number of possible miRNA seed sequences (in this case 4096 hexamers), an assumption we believe to be generally true biologically (24). Therefore, we selected MixMir6 to represent the mixed linear model results in comparison with the other methods in our analysis.

**Figure 1.**
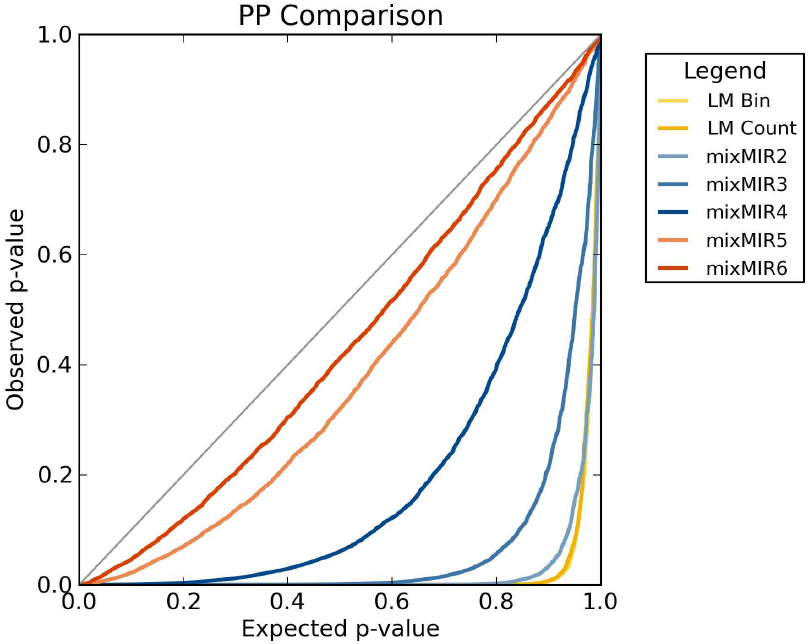
PP plot comparing the performance of MixMir with five different values of the kmer length *k* which defines the way in which the relationship matrix was constructed. The observed *p*-values are found on the *y* axis and the expected *p*-values are found on the x axis. When *p* values are correctly estimated, we would expect the observed and expected *p* values to be similar, thus approaching the *y=x* line. Here LM Bin represents the categorical linear model, and MixMir*k* represents results from MixMir with kinship matrix estimated using *k*mers. We found that higher values of *k* were better at correcting for skewness in the PP plots (i.e. false positive predictions).

The plots above assume that the *p*-values returned by the mixed linear model are properly estimated. To verify this, we performed 20 randomizations of the sequence and expression data, and ran MixMir on the randomized data for *k* = 2.6. *p*-values obtained from the randomized data exactly followed the diagonal line expected under the null hypothesis line, suggesting that our *p*-values are indeed properly estimated.

We initially tested *k* = 2 to 6 for cWords to perform model selection as we did with MixMir, which we refer to as cWords2 to cWords6. The authors recommended setting k = 1 to 3 for cWords, presumably because of the limited amount of sequence in 3’ UTRs (Methods). The resulting PP plots showed that there was a significant discrepancy between observed and expected *p*-values, similar to the simple linear models, suggesting a relatively high false positive rate for cWords on this data set, or perhaps that *p*-values are incorrectly estimated. To first test this possibility, we also performed 20 randomizations of the data as above for use with cWords. We found that the randomized data resulted in observed *p*-values again nearly identical to their expected values. Little improvement was gained by using any *k*mer background correction as judged by P-P plots (Figure S1), so we did not use this as a criterion for model selection, and relied instead on prediction performance on the T conv data set. We saw a large drop in performance for *k*=5 and *k*=6, with many fewer matches to miRBase miRNAs and T conv cell highly expressed miRNAs than for *k*=2, 3, and 4. This is entirely consistent with the authors’ recommendation and our observation above that there is insufficient 3’ UTR sequence data to train higher orders of the Markov model. Thus, for further analyses, we retained just cWords2 as representative of the algorithm.

### MixMir had the highest accuracy according to ROC curves on the T conv data

To compare MixMir against the previous motif discovery methods, we tested a total of five models: the simple linear model based on motif presence/absence (LM Bin), which we take as our baseline method, miReduce, cWords2, and Sylamer, and MixMir6. For the linear models, all possible motifs were ranked by p-value; for miReduce, we set the p-value cutoff to be 0.5, resulting in 57 motifs returned (see Methods for a discussion of this choice of p-value cutoff). Sylarray returned 885 words with p-value < 0.01, so these were ranked according to p-value. The motifs in the cWords results are ranked according to a Z-score (6), which we found were not consistent with the p-value, so we retained the original Z-score ranking, which produced better results.

We compared the significant hexamer motifs found by each method to miRNAs in miRBase (Methods). We performed two matching procedures to the miRNAs. First, in our stringent matching criterion, we considered a hexamer a match to a particular miRNA only if it matches the seed sequence of a mature miRNA. Second, in our relaxed matching criterion, we allowed the hexamers to match to any of three positions starting at nucleotides 1, 2, or 3 from the 5′ end of the mature miRNA. We included offset match positions 1 and 3 in order to include all possible types of marginal binding site matches (1), including the potential for extensive complementarity through nts 1-8. This relaxed criterion also allows for shifts in the discovered motifs, which are common in practical applications of motif-finding algorithms to biological data. In general we expect to see more false positives when including matches to offset seed sequences, so for all comparisons we considered both the results from the stringent and the relaxed matching criterion. Additional A1 type site matches (i.e. a match in the first position to an A instead of the complementary base) are discussed in the Supplementary Note.

We present results for the two matching criteria using truncated receiver operating characteristic (ROC) curves and analyze the results by computing an area-under-the-curve (AUC) value for each curve (Figure 2). Briefly, we constructed the ROC curves by taking the top 20 and 50 ranked motifs of each method, with true positives taken to be matches to any miRNA in miRBase (see Supplementary Note for details). These truncated ROC curves are exactly a close-up of the bottom left hand corner of an ROC curve over all possible results. We chose to truncate the full ROC curve, which is typically constructed over all possible 6-mer motifs, both because the methods did not return the same number of predictions and most importantly because we believe that focusing attention on only the top motifs is a more biologically meaningful comparison since only a few motifs are likely to be biologically relevant (i.e. only a small fraction of all possible miRNAs in the database are actually expressed in a cell (24)). It is important both that truncating the ROC curve does not change the ranking of the methods and that we believe our results are robust in that the ROC curves for MixMir dominate the other curves over essentially the entire range of sensitivity settings (Figure 2). We caution that the truncated AUC value should not be interpreted as a typical AUC with a baseline value of 0.5 for a random method. Instead, we plot in our curves a baseline expected value for a random predictor given the number of motifs being plotted and the number of possible true positives, to which the other AUC values may be compared.

**Figure 2.**
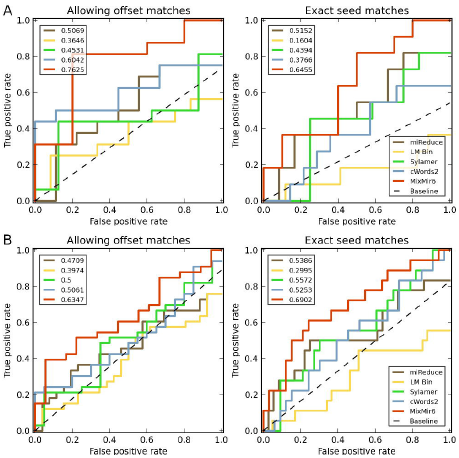
Truncated ROC curves comparing the six methods examined and their performance in ranking motifs from miRNAs in miRBase. Dotted line represents expected random performance. Top left number indicates the area under the curve for each method. Left: Results when allowing for offset seed sequence matching, Right: Results when restricting to exact seed matches only. a) ROC curve for the top 20 motifs returned by each method. b) ROC curve for the top 50 motifs returned by each method.

We present both truncated ROC curves for relaxed motif matching as well as for stringent motif matching (Figure 2). We found that the AUC values for the simple linear model was low and it found fewer miRNAs than miReduce, Sylamer, cWords2, and MixMir6. cWords2, Sylamer, and miReduce were comparable in performance in both window sizes of *N* = 20 and *N* = 50 foremost motifs. All four of those methods performed worse than MixMir6, which was more accurate over almost the entire range of sensitivity values. This effect was more noticeable when we used the strict motif matching criterion to position 2 only in the top 50 motifs. These results suggest that MixMir more accurately identifies motifs corresponding to the exact miRNA seed region. The top 50 motifs and whether they match to miRNAs in miRBase are given in Table S4.

### Validation of our computational results using experimental data sets of miRNA expression

As discussed above, one issue with the above analysis is that miRNAs are generally tissue-specific (32), and so comparing the predicted motifs to all miRNAs in miRBase, while informative, may not be the most biologically meaningful representation of their performance. We therefore further validated our results using miRNA expression levels in CD4+ T cells determined in two independent experiments by Cobb et al. (22) and Sommers et al. (23). These two experiments were conducted using different technologies, the latter measuring miRNA expression using the nCounter system (Nanostring Technologies). The Cobb et al. data compared miRNA expression profiles between CD4+CD25- and CD4+CD25+ T-cells. This experiment has the benefit of being performed in the same laboratory and on the same wildtype T conv cells from which we obtained the mRNA microarray data used in our analysis

Several, but not all, of the most highly-expressed miRNAs in each of the two data sets overlapped. Notably, the let-7 family (consisting of let-7b, let-7c, and let-7d), miR-30b, miR-26b, miR-142-3p, and miR-15a are among the miRNAs found to be expressed in T conv cells in both data sets. This is consistent with differences between the studies, including the particular labs, quantification technologies and the comparison between two cell types in the case of the Cobb et al. data.

We considered the top 20 highly expressed miRNAs found in Cobb et al. and the top 21 miRNAs found in Sommers et al. (Methods). These results can be found in Table S3. In general, we found that while there was clearly a significant overlap between highly expressed and active miRNAs, relatively few of the highly expressed miRNAs were also found to be active by the methods we tested. For example, of the top ten motifs returned, MixMir identified three exact seed sequences corresponding to highly expressed miRNAs. miReduce also performed well, but not as accurately (Tables S3 and S4). cWords found more highly expressed miRNAs than the simple linear model, but they are ranked further down the list than either miReduce or MixMir6. Of the discovered highly-expressed miRNAs, miR-142 (both 3p and 5p) is particularly interesting as it has previously been found to be highly expressed in T conv cells and it plays a significant biologically role in regulating cAMP (33). miR-142-3p was found in the Cobb et al. data and was discovered by both miReduce and MixMir6. These results suggest that miRNA motif finding algorithms can play a significant role in identifying the most biologically active miRNAs in a sample and that simply measuring miRNA expression levels is insufficient to do so.

Overall, MixMir ranked true motifs higher than other methods, while the simple linear model and cWords found fewer matches to miRNAs expressed in this cell type (Tables S3 and S4). These results are consistent with our previous analysis of the ROC curves on the full miRBase miRNA data set. These results suggest that MixMir tends to rank true miRNAs higher than other motif-finding methods, an important consideration for experimental groups that might only have the resources to validate a few top candidate miRNAs. It also shows that we were able to discover biologically meaningful results in our mouse Dicer-knockout T conv data set.

### MixMir and miReduce correct for AU bias in the motifs discovered

It is known that there is often an AU bias in computationally discovered motifs when using microarray data (34). The AU content in the 3′ UTRs used in our analyses was 55.9%, while the average AU content in the miRNA seed sequences from miRBase was 48.8%. However, the motifs discovered by the simple linear model had very high average AU content, suggesting that their high false positive rate was partially due to discovering elements representing the AU-rich background sequence (Table 1). MixMir6 motifs had average AU content similar to that in the background 3’ UTR sequence, suggesting that the correlation matrix component of MixMir successfully corrected for the AU bias. Consistent with this idea, as we altered the correlation matrix used in MixMir from *k = 2* to *k = 6*, we observed a linear decrease in the average AU content of motifs as *k* increases (see Table S5). Sylamer showed a similar degree of correction. The miReduce results had an even lower average AU content than the background 3′ UTRs. cWords, on the other hand, had motif AU composition similar to that of the simple linear model, which was very high and was not significantly changed by altering the value of *k* (Table S5). Taken together these results showed that the simple linear model suffered from high AU bias, but this bias was corrected by miReduce, Sylamer, and MixMir. Although miReduce does not have an explicit correction for 3’ UTR base composition, it likely implicitly performs this correction by finding a motif highly correlated with background composition and then finding the residuals with respect to that motif to identify the remaining motifs. We observed this phenomenon in our data in practice, where miReduce often found an AU-rich motif as the most significant motif. As described in (Methods), Sylamer likely removes the AU rich motifs as a separate preprocessing step, unlike the other methods. We discuss the possible reasons for AU bias in the Discussion section.

### MixMir corrects for 3’ UTR length and the discovered motifs are enriched for positive effects

We expect the coefficient of the fixed effect (i.e. the motif effect) to be positive if the motif represents the seed sequence of an active miRNA since miRNAs almost always downregulate their targets and a positive effect corresponds to higher expression in the Dicer KO. To test this, we looked at the number of motifs with a positive effect in each method, both overall and also compared to all motifs with a significant *p*-value (*p* < 0.01). We find that this was overwhelmingly true across all motifs, particularly the simple linear model and cWords. Sylamer returned the lowest number of positive-effect motifs in those which are significant, at only 58.64%, while MixMir showed the best enrichment for positive-effect motifs in those which are found significant, compared to the number positive over all motifs tested (Table 2).

**Table 2.**
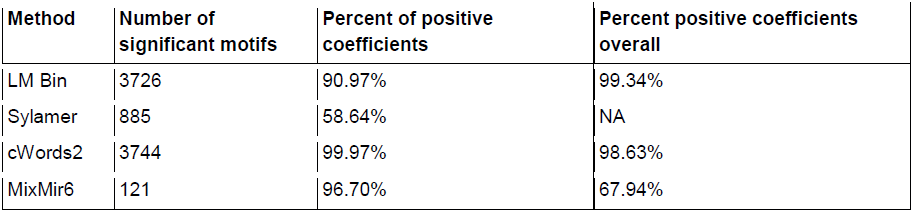
Percentage of significant motifs that have positive coefficients in the linear model. The number of significant motifs in the first column is determined by a cutoff of *p* < 0.01. The percentage of motifs from the first column which are positive (i.e., the percentage of significant coefficients which are positive) is given in the second column. The third column is the percentage of all motifs which have positive coefficients, not limited to those which have been found to be significant.

We reasoned that the overall very high enrichment of positive effects across all motifs in the simple linear model might be an artifact due to the inherent relationship between 3′ UTR length and motif count, because longer sequences have a higher probability of containing any given motif, simply by chance. Thus an mRNA that is repressed due to a miRNA motif would also induce a similar correlation for all other motifs found in that 3′ UTR. To test this hypothesis, we included 3′ UTR length as a covariate to test how it would affect the direction of the miRNA effect. A full discussion of this the 3’ UTR length effect can be found in the Supplementary Note. Briefly, the 3’ UTR length covariate strongly shifted the *p*-values of motifs found by the simple linear models, which resulted in the PP plots for the simple linear models being significantly less skewed (Supplementary Material). These results suggest that an additional reason for the higher performance of MixMir compared to the simple linear models is that MixMir implicitly corrects for 3’ UTR length using the relatedness matrix. After correcting for 3’ UTR length, we found that the percentage of positive effects across motifs remained high but not artificially high. This is consistent with our biological intuition that while most significant motifs should have positive effects, some significant motifs will appear to have negative effects due to the indirect effects that are not captured by our steady-state microarray expression measurements. In any case, since we found that the additional length covariate did not change the rankings of the top 50 motifs in any of the linear methods, we did not use it for the comparisons between methods presented above.

### All methods perform well for most miRNA transfections but MixMir performs the best for let-7b

In addition to testing MixMir on our mouse Dicer-knockout T conv data, we also tested our algorithm on miRNA transfection data from human cell lines, to demonstrate that our results are not particular to the mouse microarray data set. We tested both microarray and quantitative protein expression data obtained from Selbach et al. (20) (see Methods), and compared our results to those obtained from the same data using miReduce, cWords, Sylamer, and the simple linear model (Table 3). This data extends our analysis to a very different technology, from microarrays to pSILAC, and from mouse to human.

**Table 3.**
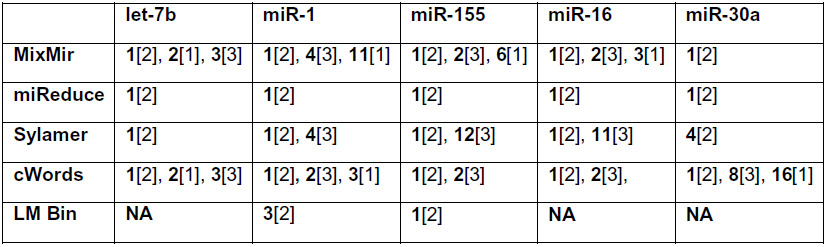
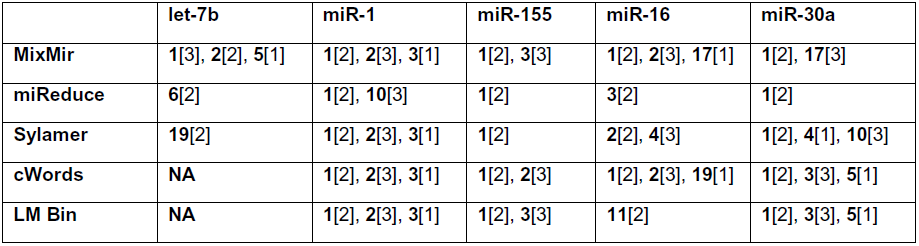
Comparison of MixMir with all other methods in miRNA transfection experiments, for (a) pSILAC quantitative proteomics, and (b) mRNA microarray data. For columns two and three, the first number is the rank of the true miRNA seed sequence, and the number in parentheses is the match position, i.e. “2” represents an exact seed match, while “1” and “3” represent offset matches. We only show results for the top 20 motifs. “NA” indicates that the true motif was not found within the top 20 motifs. miReduce was run with a *p*-value cutoff of 0.5.

We found that nearly all methods were able to find the exact seed sequence for nearly all the of the quantitative proteomics data sets, with the exception being that Sylamer ranked the seed sequence of miR-30a fourth rather than first. This is an expected result because unlike the Dicer-knockout scenario where many microRNAs were perturbed, the transfection experiment perturbs one microRNA very strongly and therefore is expected to produce much less noisy expression data. Here our analysis demonstrates that the performance of MixMir extends from the complicated T conv data set considered earlier to other simpler data sets as well.

In addition, we found that MixMir was able find the exact seed sequence or an offset seed sequence (in the case of let-7b) of the transfected miRNA as precisely the most significant motif for each of the microarray experiments at 32hrs post, while in several cases the other methods had difficulty doing so. In particular, the other statistical methods had difficulty identifying both seed and offset matches in the let-7b experiment. No other method was able to identify the seed or any offset matches in the let-7b experiment, with miReduce ranking the seed sixth, and Sylamer and cWords performing very poorly. We found that MixMir was able to find many offset seed matches - all 3 offset seed sequences were found generally within the top 10 motifs. Additionally, we found motifs further downstream of the miRNA seed sequence for let-7b (rank 17, miRNA nts 12-17), miR-155 (rank 5, nts 4-9), and miR-16 (rank 16, nts 9-14), which may be suggestive of noncanonical binding in these miRNAs (35,36). The center of miR-16 has also been suggested to be involved in binding to AU-rich elements (37) although this result has been challenged (38). Since miReduce is a useful tool in experimental labs for validating that a transfection experiment actually worked and MixMir improves on the other methods slightly for several experiments, this is an additional practical use of MixMir as well.

### MixMir predicts the miR-290 cluster as biologically most significant in mouse embryonic stem cells

Next we analyzed new unpublished microarray data from mouse Dicer knockout embryonic stem cells (Methods). We found that all methods implicated the exact seed of the miR-290 cluster (AAGTGC) as the top motif, except Sylamer which ranked it second (Table 4). It is known that the miR-290 cluster, consisting of miR-290 to miR-295, has very high activity in mouse embryonic stem (ES) cells, to the extent that replacing only this microRNA cluster can rescue most of the Dicer KO phenotype (24). Thus our results are consistent with our results for the single microRNA transfection experiments that on relatively simple experiments where only a single microRNA dominates the microRNA transcriptome of the cell, many methods are generally able to find the correct motif. However, the motif analyses extend to offset seeds and non-canonical miRNA targeting as well. MixMir was able to identify both offset seeds for the miR-290 cluster in the top ten predictions, while the other programs found either 1 or 0 of the offset seeds. We did not observe any obvious non-canonical miRNA seeds among the MixMir motifs, a point we discuss further below (Discussion).

**Table 4.** Rank of the exact seed and offset seeds of the miR-290 cluster of miRNAs for each of the methods tested for microarray data obtained from comparing Dicer knockout and WT embryonic stem cells.

|  | Microarray |
| --- | --- |
| **MixMir6** | 1[2], 6[1], 7[3] |
| **miReduce** | 1[2] |
| **Sylamer** | 2[2] |
| **cWords2** | 1[2], 6[3], 15[1] |
| **LM Bin** | 1[2] |

### MixMir identifies highly expressed miRNAs in mouse Dicer knockout adrenal cortex samples

We further tested MixMir on a published set of adrenal cortex Dicer KO experiments, performed by Krill et al. (25). The authors found that while mouse embryos with Dicer KO adrenal cortex cells developed normally up to E14.5, at E18.5 they experienced total adrenal cortex failure. In all they found 16 miRNAs that were down-regulated in the adrenal cortex of both E15.5 and E16.5 mice, including miR-34c, miR-21, miR-10a, and let-7d, which play a role in tumorigenesis among other functions (25). They also presented lists of miRNAs specifically down-regulated at each stage.

We analyzed the mRNA microarray expression data (see Methods) from both E15.5 and E16.5 embryos using the linear model, miReduce, Sylamer, cWords, and MixMir. When compared to the miRNAs that are down-regulated at both E15.5 and E16.5, we found that most methods were able to find either an exact or offset seed match to let-7d either as the first or second motif returned, with the exception of the linear model, which performed worse. Overall, MixMir ranked true miRNA seeds higher than the other methods in both E15.5 and E16.5 data sets (Table 5). Most notably, MixMir found both miR-34b and miR-34c in the top ranked motifs at E15.5, which no other method was able to do. We also performed a separate analysis of motif ranks and miRNA matches for E15.5 and E16.5 separately, as some miRNAs were found to be significantly down-regulated at one stage and not at another—namely, there were more such miRNAs at E16.5, as expected. We found similar results in this analysis, in particular that MixMir consistently found biologically significant miRNAs, with performance comparable to miReduce for both time points. cWords and the linear model were comparable for E16.5 only (Table S8).

**Table 5.** Comparison of all methods in analyses of adrenal cortex Dicer knockout data for mouse embryos at stages E15.5 and E16.5. We present matches to miRNAs found to be experimentally down-regulated in the Dicer KO samples compared to WT in both E15.5 and E16.5 adrenal cortex samples, as reported by the authors. The top 20 motifs returned by each method were analyzed. Column labeled Rank gives the rank of the motif matched; miRNAs are preceded by the match position of the motif, with [2] indicating an exact seed match.

|  | **E15.5** |  | **E16.5** |  |
| --- | --- | --- | --- | --- |
|  | **Rank** | **miRNAs** | **Rank** | **miRNAs** |
| **MixMir** | **2** | [1]miR-34b-3p, [1]miR-34c-3p | **1** | [1]miR-34b-3p, [1]miR-34c-3p |
|  | **5** | [2]let-7d-5p, [2]miR-202-3p | **3** | [2]let-7d-5p, [2]miR-202-3p |
|  | **8** | [3]let-7d | **16** | [3]let-7d-5p |
| **miReduce** | **1** | [3]let-7d-5p | **1** | [2]let-7d-5p, [2]miR-202-3p |
| **Sylamer** | **2** | [3]let-7d-5p | **NA** |  |
|  | **10** | [2]let-7d-5p, [2]miR-202-3p |  |  |
| **cWords** | **1** | [2]let-7d-5p, [2]miR-202-3p | **3** | [2]let-7d-5p, [2]miR-202-3p |
|  | **2** | [3]let-7d-5p | **9** | [2]miR-107-3p |
| **LM Bin** | **9** | [2]let-7d-5p, [2]miR-202-3p | **3** | [1]miR-34b-3p, [1]miR-34c-3p |
|  | **13** | [3]let-7d-5p | **9** | [3]miR-193a-3p |

Thus testing the different methods on additional biological data sets confirms the improvement of MixMir over previous methods. We believe that the most important use of miRNA motif finding methods is to find a small number of miRNAs that are most important in a particular cell type for further experimental validation, since usually there are very few miRNAs that are active in a cell type (24). Therefore, we view the ability of MixMir to improve the predictions by a small number of motifs to be a significant result that is a feature of the biological properties of the miRNA system.

## DISCUSSION

In conclusion, we have presented MixMir, a novel method for microRNA (miRNA) motif discovery from sequence and gene expression data. Our method corrects for pairwise sequence similarities between 3′ UTRs that could confound a motif finding algorithm in a way that is fundamentally different from previous approaches to this problem (e.g. cWords, Sylamer). We applied MixMir to a microarray dataset from wild-type and Dicer knock-out (KO) mouse CD4+CD25- T cells (T conv cells) collected by one of the authors. Since Dicer is required for miRNA biogenesis, we expect that Dicer KO cells do not contain any miRNAs and indeed this point was validated by quantitative PCR for selected miRNAs, showing a greater than 90% decrease in the knock-out (unpublished results). We found that MixMir was more accurate in finding active miRNAs in these cells than three other similar published methods, miReduce, cWords and Sylamer, as well as a simple linear regression model we used as a baseline for comparison. We validated our computational predictions using two independent biological data sets consisting of miRNA expression measurements in this cell type quantified by either miRNA microarrays or single molecule imaging using the nCounter system (Nanostring Technologies).

Importantly we found that miRNA activity was highly but not perfectly correlated with miRNA abundance in the cells, so it is not sufficient to simply measure miRNA expression levels in a cell type to determine the miRNAs that play the largest role in shaping global gene expression in those cells. For example, as in similar analyses for transcription factors, miRNAs could be highly abundant but not highly active in repressing mRNA expression due to their sub-cellular localization or the presence of competing RNA species that could sequester the miRNAs from their mRNA targets (39). Another possibility is that miRNAs may have differential efficiency of loading into the RISC complex or of targeting mRNAs, and certain mRNAs may not be efficiently repressed by miRNAs due to the presence of either stable RNA secondary structures occluding the miRNA binding site or the binding of additional *trans*-acting factors. An interesting biological finding from our analysis is that the miRNAs that we found to be the most active in T conv cells were in fact exactly the miRNAs that were more differentially expressed between these cells and CD4+ CD25+ T cells (T reg), based on previously published data from the same cell type (22).

To confirm the performance of MixMir on additional data sets, we tested MixMir against the other methods on five miRNA transfection experiments in HeLa cells, using both microarray and pSILAC quantitative proteomics data previously published by Selbach *et al*. (20). In all transfection experiments, for the pSILAC data, nearly all methods were able to find the exact seed sequence first, with the exception of the linear model, which failed to do so in three cases, and Sylamer, which failed to do so for miR-30a (Table 3a). In the microarray data, MixMir ranked the exact seed sequence of the transfected miRNA first, with the exception of let-7b where it ranked it second. For the let-7b experiment, all of the other methods performed much more poorly than MixMir, demonstrating that MixMir gives a significant improvement on at least one transfection experiment.

We performed a similar analysis with mouse embryonic stem (ES) cell Dicer knockout experiments, for which we also included previously unpublished microarray expression data, and mouse adrenal cortex Dicer knockout experiments, using data obtained from Krill et al. (25). In the former, we found again that most methods we tested were able to identify the seed sequence of the miR-290 cluster known to be highly active in ES cells but that MixMir additionally found more offset seed sequences for this cluster; in the latter, we found that MixMir either identified more true miRNAs or performed comparably to the other methods depending on the time point examined. Note that the adrenal cortex data might be noisier than the other experiments because it was derived from more heterogeneous primary tissue rather than cell cultures.

These experiments demonstrate the general applicability of MixMir on different technologies (microarray and proteomics), species (human and mouse), cell types (cell lines, primary T cells and adrenal cortex tissue) and experiments of varying complexity, from relatively simple (microRNA transfection or ablation of a single dominant microRNA cluster) to relatively complex (perturbation of many microRNAs in a tissue). Our analysis of HeLa cells also demonstrate the utility of MixMir in a context where miReduce is often used in practice – to verify that a miRNA transfection experiment was carried out successfully.

In our miRNA targeting model, we made several assumptions similar to previous methods, like miReduce. First, we searched over non-degenerate *k*mer motifs only. Although this does not rule out the possibility of detecting degenerate motifs, it probably biases our search towards non-degenerate seed matches. Although we searched for several published types of degenerate motifs such as G-bulge sites and imperfect sites in our data, we found only a few cases of such sites. We note that many of the analyses of non-canonical miRNA motifs have been performed on Ago HITS-CLIP or PAR-CLIP data and therefore represent biochemical binding events of the miRNAs, which are not necessarily perfectly correlated with repression that is detectable at the mRNA level. Similar observations hold for ChIP-seq data on transcription factors where biochemical binding does not necessarily produce transcription of the target gene. Second, we searched over motifs in 3’ UTRs only. This choice was based on previous results in the literature but can be easily changed to examine other sequences, such as coding sequences or 5′ UTRs, by users of MixMir. Third, our model assumes that the miRNA regulatory effect is additive, which is supported by previous evidence (1) but is still an approximation to biological reality.

Our approach to the motif discovery problem borrows an idea from genome-wide association studies (GWAS), namely that cryptic relatedness between individuals acts as a confounding factor that causes simple linear models to detect many false positive associations. In GWAS, cryptic relatedness is captured by a kinship matrix representing pairwise similarities between individuals. In the miRNA motif discovery problem, we considered background nucleotide composition similarity, which may affect miRNA binding in a variety of ways. It may affect binding site accessibility (12), represent other *cis*-regulatory sites for RNA-binding proteins, or simply be a correlate of paralogy – consider for instance ribosomal genes that are very similar and have similar expression patterns (e.g. due to similar transcriptional regulation) but are not affected by miRNA targeting (40). Such signals can confound a motif finder based on a simple linear model if sequence similarity is not corrected. In particular, we found that the relatedness matrix corrected for high AU content of the 3’ UTRs. This observation could be due to the presence of AU-rich elements, which are known to be involved in mRNA regulation, other AU-rich motifs for *trans*-acting factors or more open secondary structures in the 3’ UTR that might increase the efficiency of miRNA binding.

We constructed a relatedness matrix analogous to the kinship matrix by representing *k*mer content similarity between 3′ UTRs, which implicitly accounts for 3’ UTR length. Our finding *k* = 6 provided the most correction of the results is intuitive, as this choice of *k* corrects for motifs of the same length as the seed sequences for which we are searching and synergistic interactions between nearby miRNA binding sites and RNA binding protein binding sites have been previously documented (12,41,42). It is possible that we are also computing an approximation to the alignment score of the 3’ UTRs and that global similarity of 3’ UTRs is more important than the presence of short, 6 nt motifs, but we consider this possibility unlikely because very few pairs of 3’ UTRs should have any meaningful sequence alignment at all. Most significantly, MixMir6 was able to correctly implicate significant hexamer motifs associated with both known miRNAs as well as with highly expressed miRNAs in our dataset, as indicated using the area under the truncated receiver operating characteristic (AUROC). In particular, on the data sets we tested, MixMir performed better than current state-of-the-art methods of motif discovery, miReduce, cWords, and Sylamer (5–7) over the entire range of sensitivity settings considered and on several different types of data.

Notably, both cWords and Sylamer correct for 3′ UTR length and compositional biases. Overall, cWords performed better than Sylamer, but exhibited strong AU bias in the datasets we examined. These results suggest that background nucleotide composition similarity can strongly affect the ability of a linear model to uncover true motifs, but also that the way in which we correct for background composition can dramatically alter the results. Unlike Sylamer and cWords, MixMir utilizes the expression fold change values instead of just the ranks. Additionally, MixMir makes pairwise comparisons of the entire 3′ UTR sequences, thus performing a more direct comparison of sequence context, rather than comparing motif vs. background composition within each 3′ UTR like the other methods.

The MixMir software is freely available online, and utilizes FaST-LMM, a fast mixed linear model solver that can be obtained freely online (see the MixMir README file with the software for details). One limiting factor in our approach is the total amount of 3’ UTR sequence available to construct large correlation matrices for long kmers. Increasing the kmer length to 7mers or higher would make the correlation matrix very sparse and difficult to estimate accurately. Another drawback of MixMir is its relative computational inefficiency: we exhaustively analyzed all 6mers but if we wanted to exhaustively analyze all 7-or 8mers, the runtime and memory requirements for FaST-LMM would make the computation too inefficient for practical use in our experience. However, miReduce suffers from a similar problem of computational inefficiency for values of k greater than about 6, so this is not an issue unique to MixMir.

Finally, we note that our mixed linear model approach is not limited to solving the miRNA motif discovery problem. Like the REDUCE software, MixMir can potentially also be applied to other regulatory element motif detection problems, such as transcription factor and RNA binding protein motif prediction, by varying the type of sequence input and gene expression fold change input. For example, REDUCE was originally applied to transcription factors but was later applied to miRNAs in miReduce (7), RNA binding proteins in matrixREDUCE (43), degenerate transcription factor motifs in fREDUCE (44) and ChIP-chip data. We believe that MixMir can be similarly applied to many of these types of data and possibly also other data types such PAR-clip (45) as well.

## ACKNOWLEDGEMENT

We thank Matthias Merkenschlager for providing laboratory space for producing the experimental data used in this study and helpful discussions. We also thank Marc Friedlaender, Dominic Gruen, Alexander Schliep, Michael Seiler and Jinchuan Xing for comments on this work.

